# CRISPR-Cas13 mediated Knock Down in *Drosophila* cultured cells

**DOI:** 10.1101/2020.11.01.364166

**Authors:** Raghuvir Viswanatha, Michela Zaffagni, Jonathan Zirin, Norbert Perrimon, Sebastian Kadener

## Abstract

Manipulation of gene expression is one of the best approaches for studying gene function *in vivo*. CRISPR-Cas13 has the potential to be a powerful technique for manipulating RNA expression in diverse animal systems *in vivo*, including *Drosophila melanogaster*. Studies using Cas13 in mammalian cell lines for gene knockdown showed increased on-target efficiency and decreased off-targeting relative to RNAi. Moreover, catalytically inactive Cas13 fusions can be used to image RNA molecules, install precise changes to the epitranscriptome, or alter splicing. However, recent studies have suggested that there may be limitations to the deployment of these tools in *Drosophila*, so further optimization of the system is required. Here, we report a new set of PspCas13b and RfxCas13d expression constructs and use these reagents to successfully knockdown both reporter and endogenous transcripts in *Drosophila* cells. As toxicity issues have been reported with high level of Cas13, we effectively decreased PspCas13b expression without impairing its function by tuning down translation. Furthermore, we altered the spatial activity of both PspCas13b and RfxCas13d by introducing Nuclear Exportation Sequences (NES) and Nuclear Localization Sequences (NLS) while maintaining activity. Finally, we generated a stable cell line expressing RfxCas13d under the inducible metallothionein promoter, establishing a useful tool for high-throughput genetic screening. Thus, we report new reagents for performing RNA CRISPR-Cas13 experiments in *Drosophila*, providing additional Cas13 expression constructs that retain activity.

## Introduction

One of the most powerful ways to investigate gene function is to modulate the expression of specific genes. This approach is even more informative if the perturbation can be done in a time and a tissue-specific manner. *Drosophila* has been an excellent model for this type of genetic-phenotypic approaches over the years, mainly using the GAL4-UAS system, that allows controlled expression of transgenes^1^. For example, the levels of specific mRNAs can be modulated by expressing short hairpin RNAs (shRNA) or overexpression constructs in a time and space-controlled manner *in vivo*^2^. However, the use of shRNAs for knocking down (KD) genes has several problems^3,4^. First, the knockdown is not always complete and some cells (i.e. neurons) seems to be particularly resistant to shRNAs. Second, expression of different shRNAs against the same transcript often leads to inconsistent phenotypes, raising the concern of possible off-target effects that could result in misleading conclusions. Indeed, off-targets effects are often difficult to identify, despite the use of proper controls, such as the expression of shRNA resistant transcripts to rescue the phenotype.

CRISPR and CRISPR-like systems were originally discovered as adaptative immune effectors in prokaryotes and archaea that degrade foreign nucleic acids^5^. The Type-II CRISPR systems are composed of two elements: a guide RNA (crRNA) and an endonuclease. The crRNA consists of a constant stem loop and the reverse complement sequence of the region to be targeted. The crRNA associates with the Cas enzyme and directs it to the target sequence. The discovery that Cas enzymes and artificial crRNAs can be expressed in eukaryotic models to target endogenous sequences has been a major technological breakthrough^6–8^. Indeed, DNA targeting CRISPR enzymes, such as Cas9 and Cas12, are now commonly used for genome editing and genetic screenings in cell culture and *in vivo*. Moreover, the same system can be used to activate the expression of specific genes from their endogenous locus by using crRNAs to target enzymatically-inactive (“dead”) Cas9-VP16 fusion proteins to the desired gene. This technology has been successfully adapted to *Drosophila* and provides a universal way to overexpress genes avoiding artifacts derived from cDNA overexpression^9^. However, Cas9-based approaches such as (“dead”) Cas9-repressor^10,11^ have failed until now to generate a tool to downregulate gene expression in *Drosophila* (unpublished observations).

Recently, CRISPR-Cas13 has been characterized and used to achieve mRNA knock down and manipulation in mammalian cells^12–14^. Indeed, Cas13 is so far the only known family of Type-II CRISPR-Cas enzymes that has RNAse activity, provided by the presence of Higher Eukaryotes and Prokaryotes Nucleotide (HEPN)-binding domains^15,16^. Cas13 forms complexes with the ∼64 nt long crRNAs by recognizing a particular stem loop structure. Then, the Cas13-crRNA complex identifies its target through the ∼30 nt long protospacer present in the crRNA^16^. To date, four Cas13 subtypes (a, b, c, and d) have been identified^17^ and categorized according to the differences in both protein and crRNA stem loop structures^16,18^. Although all the Cas13 subtypes can be used for targeted mRNA knock down in mammalian cells, PspCas13b and RfxCas13d are the most efficient, providing up to 90% expression reduction of a reporter gene^13,14,19^. Importantly, in bacteria, Cas13 displays collateral non-specific RNase activity of single stranded RNA that are in proximity to the enzyme^20^. However, this phenomenon has not been observed in mammalian cells, making Cas13 an amenable tool for targeted knockdown that could be utilized as an alternative to RNAi or a validation of this approach.

Unfortunately, expression of RfxCas13d is linked to toxicity *in vivo*^21,22^, raising concerns about their possible applications. For example, Buchman et al., showed that the ubiquitous co-expression of Cas13d and a crRNA to target either *white (w), Notch (N)*, or *yellow (y)* mRNAs is toxic and leads to lethality. Therefore, the use of Cas13 in *Drosophila* requires further optimization for reducing toxicity and providing a reliable tool. Here, we generated and tested the functionality of several new PspCas13b and RfxCas13d constructs in S2 cells. Moreover, we generated a versatile experimental toolbox that allows tight control of the expression and localization of Cas13. Importantly, expression of the Cas13 variants along the crRNA provide a robust silencing power (70% KD efficiency) of both reporter and endogenous transcripts. Furthermore, we established a new S2 cell line that stably expresses RfxCas13d that can be used for high throughput genetic screening.

## Results

### Cas13 efficiently downregulates transfected reporter transcripts in S2 cells

To determine whether the two “next generation” Cas13 isoforms, PspCa13b (referred to as Cas13b) and RfxCas13d (referred to as Cas13d)^13,14,19^ can be used to specifically downregulate mRNAs, we tested their abilities to downregulate the activity of transfected *luciferase* and *mCherry* reporter transgenes in *Drosophila* S2 cells. Briefly, we co-transfected a plasmid containing a UAS-driven *PspCas13b*, a constitutive *Actin*-promoter-driven Gal4 (pActinGal4), a copper-inducible luciferase reporter, and a guide RNA complementary to the luciferase coding sequence into *Drosophila* S2 cells. To diminish nuclear accumulation of the Cas13 protein, we added a Nuclear Exportation Signal (NES) to the PspCas13b. In addition, the expressed Cas13b protein contained three tandem HA tags (3XHA). Following transfection and copper induction, we obtained a 70% and 46% signal reduction of Renilla or Firefly luciferase activity, respectively, in the Cas13b + crRNA vs control cells (Fig. 1A, B). Importantly, the decrease of luciferase activity is dependent on the catalytic activity of PspCas13b, as a catalytically inactive version of Cas13b (dCas13b) was unable to reduce luciferase activity (Fig. 1A, B). Furthermore, and similar to reported in mammalian cells^14,19^, the expression of the crRNA alone did not result in lower levels of the reporters, demonstrating that the crRNA does not act though the RNAi pathway. Thus, these results demonstrate that PspCas13b provides specific knockdown of a reporter gene in *Drosophila*. The differences in the knockdown efficiencies are likely due to the design of the crRNA (see below).

**Figure 1:**
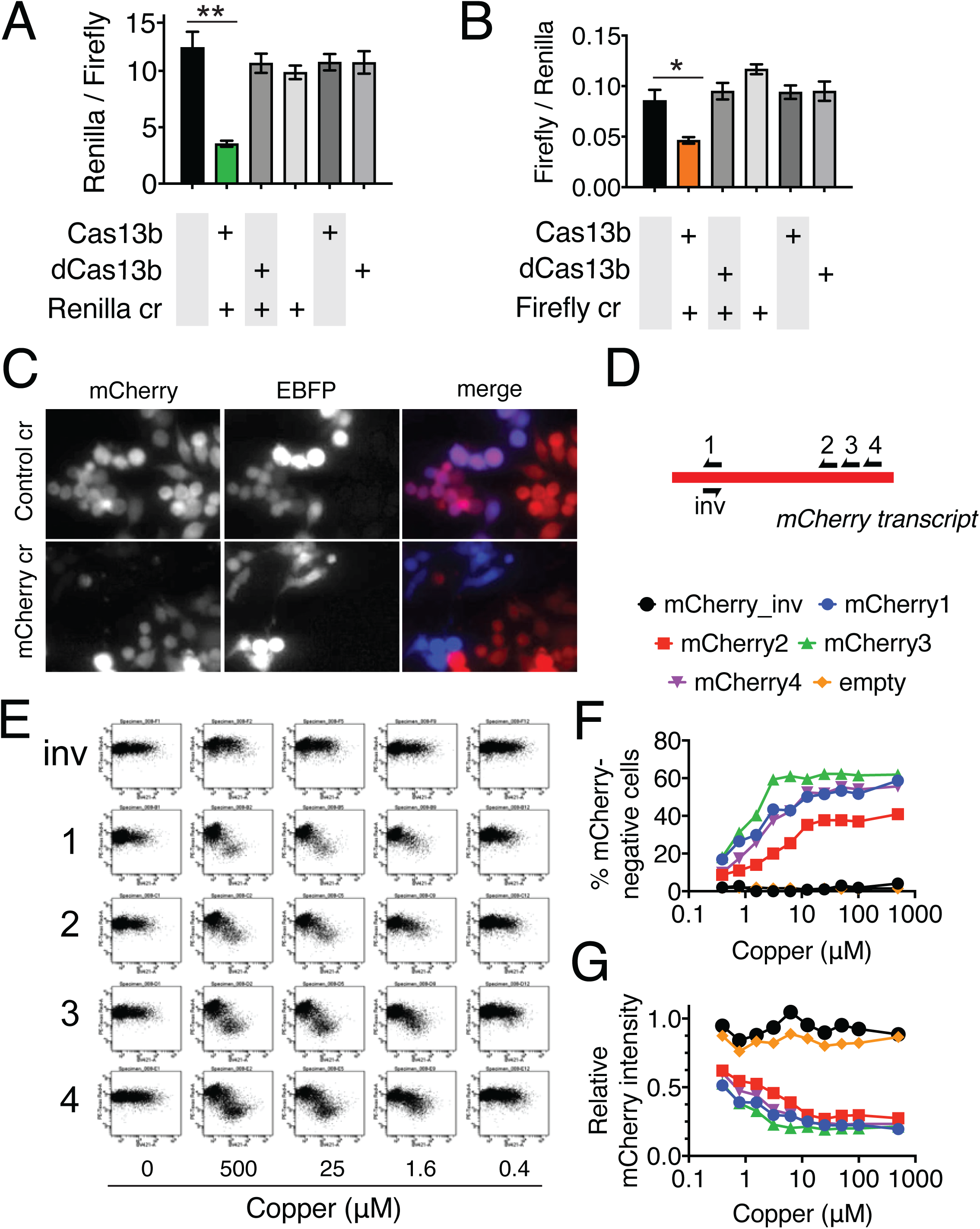
Cas13 can mediate efficient knockdown of reporters in *Drosophila* cells. A) Luciferase assay showing KD of Renilla luciferase by Cas13b (*UAS-PspCas13b-NES-3XHA*) in S2 cells. Error bars represent the SEM of 3 independent biological replicates. Two-tailed t-test, ** p-value = 0.004, p-value >0.05 if not indicated. Cells were induced with 500μM copper sulfate to allow the expression of luciferase 48 hours after transfection. The luciferase assay was performed 24 hours after the induction (see methods). B) Luciferase assay showing KD of Firefly luciferase by Cas13b in S2 cells. Error bars represent the SEM of 3 independent biological replicates. Two-tailed t-test, * p-value = 0.0275, p-value >0.05 if not indicated. We applied the same protocol described for Fig. 1A. C) Image showing KD of endogenous mCherry by Cas13d (stably transfected, copper inducible MT-NLS-RfxCas13d-NLS-HA, induced by 100 µM copper for 4 days). The plasmids expressing crRNAs co-express EBFP. D) Schematic representing locations and direction of the crRNAs along the mCherry transcript. E) Flow cytometry plots of mCherry fluorescence (*y*-axis) versus EBFP intensity (*x*-axis) for cells expressing indicated crRNA under indicated copper induction. F) Quantification of number of EBPF-positive cells with low mCherry fluorescence. G) Quantification of average mCherry intensity of EBFP-positive cells.

### Cas13 efficiently downregulates an endogenous reporter transcript in S2 cells

We next tested whether the CRISPR-Cas13 system can knockdown an endogenously-expressed mCherry fusion protein. For this experiment we first generated a stable cell-line expressing copper-inducible metallothionein (MT) promoter-driven Cas13d (pMT-NLS-Cas13d-NLS-HA-T2A-GFP) in NPT005 cells, in which *mCherry* is fused to the *Clic* gene via an intronic insertion^23^. By screening sublines from this stable transfection, we isolated and expanded a colony with strictly inducible GFP expression, suggesting that Cas13d activity would also be inducible. We then transiently transfected these cells with a crRNA complementary to a region of the *mCherry* coding sequence or a control crRNA, and then induced Cas13d expression. The crRNA vector co-expresses actin promoter-driven *EBFP* as a marker of transfection level. The results demonstrated significant reduction of the endogenously expressed mCherry reporter in a *mCherry* crRNA-dependent manner (Fig. 1C).

### Cas13 enables programmable gene titration

Some proteins show dose-dependency in controlling certain biological process and in these cases, gene titration is a necessary approach for comprehensively study their functions. The CRISPR-Cas13 system relies on an entirely exogenous machinery and thus is expected to be titratable. To verify whether knockdown could be titrated by changing Cas13d induction levels in *Drosophila* cells across various crRNAs, we first chose 4 regions along the *mCherry* coding sequence and created crRNAs complementary to these regions or an inverted (sense) control for region 1 (Fig. 1D). We transiently transfected plasmids expressing these crRNA expression (and co-expressing *EBFP*) in MT-Cas13d cells induced with varying copper levels. To determine the relationship between crRNA amount (as indicated by EBFP; *x*-axis) and target mRNA level quantitatively (as indicated by mCherry; *y*-axis), we subjected cells to flow cytometry (Fig. 1E-G). The results showed that we can achieve copper-dependent gene titration with any of the chosen *mCherry* complementary crRNAs but not with the inverted crRNA (Fig. 1E). We gated the cells to account for incomplete transfection efficiency (Supplementary Fig. 1). This revealed that Cas13d level dramatically affected both the fraction of crRNA-expressing cells that were mCherry-negative and the overall intensity of mCherry among crRNA-expressing cells (Fig. 1F,G). We did not observed any change in the percentage of mCherry positive cells or in mCherry signal-per-cell for each tested crRNA when Cas13d expression was not induced, indicating that the system is not leaky in this cell-line and that the crRNA does not trigger RNAi by pairing to the target transcript as shown in Fig 1A,B. Thus, Cas13 is amenable to highly specific, titratable knockdown in *Drosophila*. Taken together, our data demonstrate that both Cas13b and Cas13d efficiently knockdown transient and endogenous reporter transcripts in S2 cells. Importantly, we showed that the system is highly tunable by regulating the amount of the Cas13 effector and that the crRNA per se does not trigger RNAi.

### CRISRP-Cas13 knockdown is robust to changes in localization, variant, and expression level

Next, we evaluated a toolbox of *Drosophila* Cas13 constructs in which we changed the subcellular localization, variant, or expression level and measured knockdown of the endogenous *mCherry* reporter. We constructed equivalent UAS-driven Cas13b or Cas13d variants fused to a strong nuclear export signal (NES) or tandem nuclear import signals (NLS) and expressed these in *mCherry* reporter cells along with crRNAs complementary to *mCherry* or an inverted control (Fig. 1D). Our Flow Cytometry analysis showed that for each tested crRNA, there is not significant difference between *UAS-RfxCas13d-NES-3XHA* and *UAS-NLS-RfxCas13d-NLS-3XHA* or between *UAS-PspCas13b-NES-3XHA and UAS-NLS-PspCas13b-NLS-3XHA* (t-test, p > 0.05), indicating that the subcellular localization of Cas13 does not alter its knockdown efficiency for the tested mRNA (Fig. 2A; Supplementary Fig. 2A). Then, we compared PspCas13b activity with RfxCas13d. We observed similar knockdown efficiency for RfxCas13d and PspCas13b when co-expressed with *mCherry1* and *mCherry2* crRNAs. However, RfxCas13d showed a diminished knockdown efficiency than PspCas13b in combination with *mCherry3* and *mCherry4* crRNAs. These data indicate that optimal crRNA design might be more critical for RfxCas13d than for PspCas13b (Fig. 2A). However, when stably expressed and then induced from the MT promoter, RfxCas13d is more effective than the transiently expressed Cas13b. These results likely reflect lower levels of Cas13d when transiently transfected in *Drosophila* cells compared with Cas13b, and significant overexpression when expressed stably (data not shown).

**Figure 2:**
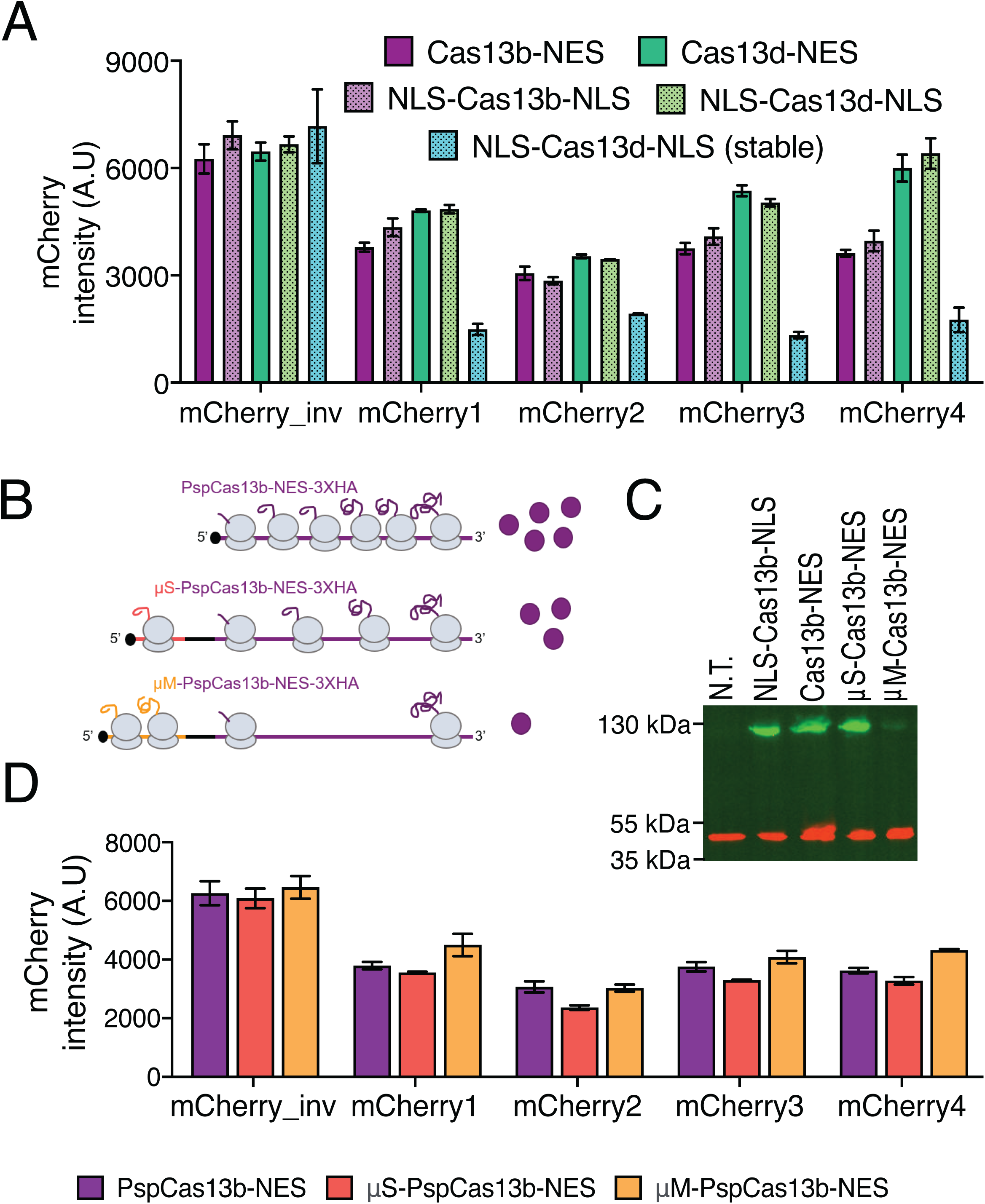
Side-by-side comparison of Cas13 variants localization, subtype, and expression. A) Flow Cytometry quantification showing the KD of mCherry in cells co-transfected with indicated Cas13 variants and crRNAs. Error bars represent the S.E.M. of 3 independent biological replicates. B) Schematic representing the variants of Cas13b with the upstream ORFs (μORFs) to tune Cas13b translation (*UAS-μS-PspCas13b-NES-3XHA, UAS-μM-PspCas13b-NES-3XHA*). The two stop codons between the μORF and the Psps-Cas13b-NES-3XHA cassette are represented by the black portion of the polycistronic mRNA, between the μORF and the PspCas13-NES-3XHA cassette. C) Western blot showing the expression of the Cas13b variants using α-HA antibody to detect the 3XHA tag. N.T. stands for “not transfected”, used as negative control. Detection of Actin was used as loading control. D) Flow Cytometry quantification showing the KD of mCherry with *UAS-PspCas13b-NES-3XHA, UAS-μS-PspCas13b-NES-3XHA*, and *UAS-μM-PspCas13b-NES-3XHA*. Inverse (inv) refers to the crRNA used as a control, carrying the sense sequence. Error bars represent the S.E.M. of 3 independent biological replicates.

As mentioned above, toxicity of Cas13 is a major barrier to its use *in vivo*^22^, and may be overcome by decreasing its expression. To modulate Cas13 effector expression using the GAL4-UAS system, we explored the use of an upstream open reading frame (μORF)^24^. This system allows tunable expression of a given open reading frame because the translational efficiency of the downstream cassette is inversely proportional to the length of the μORF. We generated plasmids carrying the PspCas13b-NES-3XHA and two different μORFs, one of 22 and another of 38 amino acids, named μS (short) and μM (medium), respectively (Fig. 2B). As expected, introduction of the longer μORF (μM) resulted in a strong decrease in the levels of PspCas13b-NES-3XHA protein, while the presence of the μS μORF did not considerably alter the amount of PspCas13b protein, as assessed by Western Blot (Fig. 2C). We then determined the capacity of the μORF-containing PspCas13b constructs to knockdown reporter transcripts. For this experiment, we used the aforementioned four different crRNAs targeting the fluorescent protein mCherry, stably expressed by the cells, and quantified the knockdown by Flow Cytometry analysis (Fig. 2D; Supplementary Fig. 2B). For each tested crRNA, the knockdown efficiency was similar among all PspCas13b-NES-3XHA variants (two-way ANOVA, p-value < 0.0001), suggesting that the μM μORF is ideal as it achieves maximum knockdown activity with minimal PspCas13b expression.

As hypothesized before, our data indicate that knockdown optimization requires the design of multiple crRNAs against the same target. In fact, the *mCherry2* crRNA appeared to be significantly more efficient than *mCherry1* and *mCherry3* (Tukey’s multiple comparisons test, p < 0.05 and p<0.5, respectively) for *UAS-μS-Psp-Cas13b-NES-3XHA* and the most efficient crRNA among the four tested for *UAS-μMPspCas13b-NES-3XHA* (Tukey’s multiple comparisons test, p < 0.005 mCherry, p< 0.5 mCherry3, p<0.05 mCherry 4).

In conclusion, these experiments showed that PspCas13b is highly effective in transient assays, since the knockdown effect is preserved despite dramatic reduction on the protein levels by the presence of the upstream ORFs. Furthermore, we showed that both PspCas13b and RfxCas13d can degrade mRNAs independently of their nuclear or cytoplasmic localization.

### RfxCas13d can be used to downregulate an endogenous mRNA target and generate an observable phenotype in *Drosophila* cells

We further tested the functionality of this system by targeting an endogenous transcript in S2 cells. For this purpose, we utilized either the Cas9 or Cas13 systems to target Rho1, a GTPase involved in cytoskeleton remodeling pathways^25,26^. Among other functions, Rho1 mediates the assembly of the actin contracting ring to drive cytokinesis. Hence, cells depleted of Rho1 fail to complete cytokinesis and, as a result, are polynucleated and have a larger than normal diameter^27^. First, we observed that the phenotype induced by Cas9 *Rho1* knockout is similar to the knockdown mediated by Cas13. As expected, co-expression of Cas9 with Cas13-crRNA along with co expression of Cas13 with a Cas9 sgRNA does not induce any observable cell enlargement (Fig. 3A). Then, we optimized the system by designing multiple gRNAs and testing their efficiency in the S2 cells stably expressing MT-NLS-RfxCas13d-NLS. We indeed observed a significant cell enlargement (cells larger than 15μm) of cells with the most efficient tested crRNA (*Rho1-1*). Importantly, the penetrance of the phenotype depended on the expression levels of NLS-RfxCas13d-NLS-HA. Indeed, for each tested crRNA, increasing concentrations of copper sulfate result in increasing proportions of enlarged cells (Fig. 3B,C), indicating that the effect is titratable in a similar range compared with the effect on mCherry. Moreover, we showed that the phenotype is not dependent on the strictly inducible cell clone or the presence of copper sulfate, as a similar phenotype is observed in non-inducible MT-NLS-RfxCas13d-NLS stable cells (clone D7, which retains leaky activity in the absence of copper sulfate) (Supplementary Fig. 3). In conclusion, our data demonstrate that Cas13 can be used to target endogenous transcripts and that the knockdown activity is tunable.

**Figure 3:**
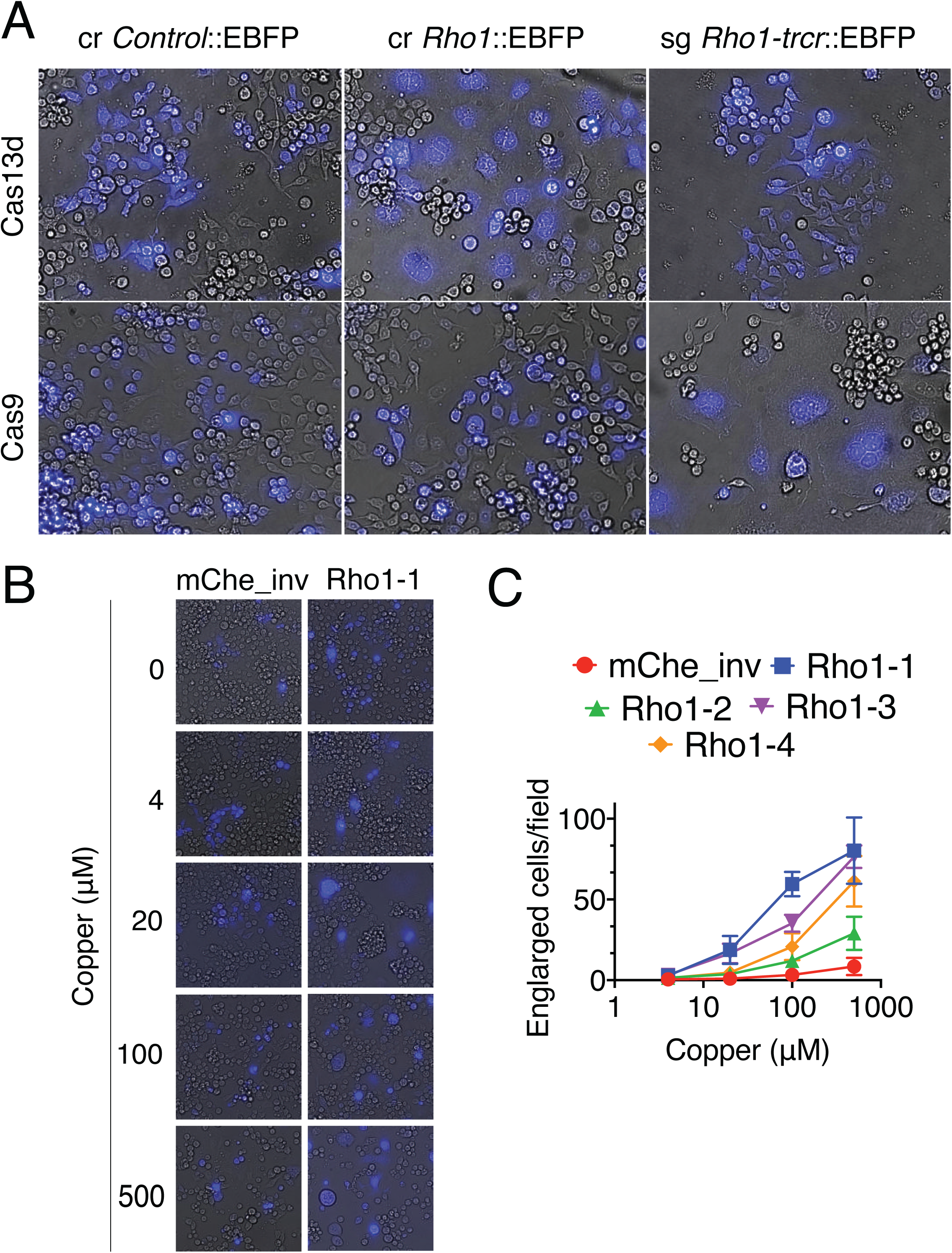
Cas13 mediated knockdown of an endogenous target causes a visible phenotype. A) Representative images showing the phenotype of cells expressing Cas13d (stably transfected, copper inducible MT-NLS-RfxCas13d-NLS-HA, induced by 100 µM copper for 4 days) or Cas9 and indicated crRNA or sgRNA. crRNA and sgRNA expression vectors co-express EBFP. B) Phenotypic series of cells expressing indicated crRNA (marked by EBFP) induced by variable copper dosage. C) Quantification of *Rho1* KD with increasing amount of copper concentrations to induce the expression of NLS-Cas13d-NLS-3XHA. As a control, we used the crRNA carrying the sense sequence of mCherry (mChe_inv). The field area is 0.44 μm^2^.

## Discussion

Here, we describe the use and generation of a new toolkit to perturb gene expression in *Drosophila*. The CRISPR-Cas13 system can be used for RNA knockdown with greater accuracy than RNAi^13,19^, but also to image RNA molecules^12,28^, manipulate the epi-transcriptome^29,30^, edit transcripts^14^, and alter splicing^19,31^, but the use of this system in *Drosophila* has been limited by the potential toxicity of the nucleases^22^. Here, we report additional PspCas13b and RfxCas13d expression constructs that successfully knockdown transcripts in *Drosophila* cells. First, we generated a new set of *UAS-PspCas13b* and *UAS-RfxCas13d* expression vectors and developed a cell-based system to evaluate their knockdown efficiency. We showed that this system can efficiently knockdown a transcriptional reporter or an endogenous gene. Second, we demonstrated that both expression and activity of Cas13 can be tuned by the use of an inducible promoter construct. Previous experiments from our group have shown that accounting for dose-dependency can quantitatively improve screen reproducibility and robustness, particularly for context-specific cell-essential genes, such as tumor cell vulnerabilities^32^. Our *Drosophila* cell-line is compatible with large-scale pooled screening, and therefore could be used to perform pooled, variable-dose screening^33^. Last but not least, we achieved tunable expression of PspCas13b by decreasing its translation through the addition of upstream ORFs of different lengths. This strategy has successfully been employed in many biological contexts, including for balancing Cas9 activity and toxicity in *Drosophila*^24^.

There is an obvious need to develop RNA CRISPR-Cas13 in *Drosophila*. First, despite the large number of RNAi-expressing *Drosophila* stocks there is still a limited number of transgenic shRNA reagents targeting any given *Drosophila* gene. Because of the significant risks of false positive and false negative results, a single shRNA line against a targeted gene is not sufficient to provide a high-confidence result. Moreover, gaps in coverage remain, as no transgenic shRNA reagents exist for ∼5000 genes and ∼6000 genes are only covered by a single shRNA reagent ^2^. Moreover, the reported improvement in targeting efficiency and decreased off-targeting may be useful to increase coverage and usefulness of the existing *Drosophila* loss-of-function reagent stock. Indeed, while the shRNA targets -21nt, the crRNA here design targets a -30nt, increasing the specificity of the system and decreasing the likelihood of off-targets effects. Second, while RNAi relies on host cell machinery, RNA CRISPR only relies on Cas13 and crRNA, and thus is easily tunable by controlling Cas13 expression, as we demonstrate. This may be useful *in vivo* to achieve various levels of knockdown depending on the developmental stage or tissue being examined and could be accomplished in one cross. Third, inactive Cas13 fusions have been used to precisely edit the epi-transcriptome or alter splicing^14,21,29–31^. Utilizing these reagents in *Drosophila* could reveal the developmental need for epi-transcriptome modification of transcripts in time and space and could be used to determine the developmental function of particular splicing changes.

In summary, we showed that RNA CRISPR-Cas13 robustly works in *Drosophila* cells for targeted mRNA knockdown. Further experiments will be needed to determine whether our Cas13 variants can overcome *in vivo* toxicity issues and be used in flies as an orthogonal approach to RNAi for whole-body or tissue-specific RNA modification.

---

**Supplementary Figure 1: Gating strategy related to Figure 1**. A) Scheme for analyzing flow cytometry data in Figure 1F,G. B) Representative flow cytometry plots and gating.

**Supplementary Figure 2**: **Representative contour plots related to Figure 2**. A) Representative flow cytometry plots for Figure 2A. B) Representative flow cytometry plots for Figure 2D.

**Supplementary Figure 3**: The **Rho1 phenotype is not due to a clonal effect or the presence of copper sulfate**. A non-inducible (constitutive) clone expressing MT-NLS-Cas13d-NLS-HA-T2A-GFP was transfected with a crRNA vector co-expressing EBFP and assessed for cytokinetic defects. Cell-enlargement was observed in transfected cells 4 days post-transfection.

## Acknowledgments

We thank the Marr lab at Brandeis University-MA for sharing S2 cells. We thank Daniela Ischiu-Gutierrez and the Harvard Immunology Flow Core for flow cytometry support. We warmly thank Nicholas Brideau of the Hsu Lab and all the members of Kadener and Perrimon lab for helpful and critical discussion. Special thanks to Dr. Irina Marcovich for her crucial advice and technical support for the cloning of PspCas13b variants.

## Abbreviations

PspCas13b: Prevotella sp. P5-125 Cas13b
RfxCas13d: Ruminococcus flavefaciens Cas13d

## Methods

### Cell culture and transfection

S2-C1 cells were a kind donation from Dr. Marr lab at Brandeis University, Waltham -MA. S2 cells were maintained in Shields and Sang M3 Insect Medium (Sigma), supplemented with 10% of Heat inactivated Fetal Bovine Serum (Genesee) and 1% Penicillin/Streptomycin (Sigma). S2 cells were transfected with Bio TransIT-2020 Reagent (Mirus), according to the manufacturer instructions (2μg of total DNA). 0.5×10^6^ S2 cells/well were plated the night before transfection into 12-well plates. S2R+ cells and S2R+-NPT005 cells were obtained from the Drosophila Resource Screening Center at Harvard Medical School and maintained in Schneider’s media (Thermo 21720), 1X Penn/Strep (Thermo 15070063), 10% Heat inactivated Fetal Bovine Serum (Thermo 16140071). S2R+, S2R+-NPT005, and S2R+-NPT005/MT-NLS-Cas13d-NLS-HA-T2A-GFP_CloneF11 cells were transfected using Effectene (Qiagen) according to manufacturer instructions. 1×10^6^ cells/well were seeded 30 minutes before transfection on 24-well plates.

### Luciferase assays

48 hr after transfection, Firefly luciferase and Renilla luciferase expression was induced with 500 μM copper sulfate for 24 hr before performing the Luciferase Assay. Briefly, cells were pellet by centrifugation and lysate in lysis buffer (25 mM Tris-phosphate at pH 7.8, 10% glycerol, 1% Triton X-100, 1 mg/ml of bovine serum albumin (BSA), 2 mM CDTA (cyclohexylenediaminetetraacetate), and 2 mM DTT). An aliquot of the lysate was added to Firefly Buffer 1X (75 mM Tris pH 8.0,100 μM EDTA, 530 μM ATP, 5mM MgSO_4_) freshly supplemented with 0.1 μM D-luciferin and 100 μM Coenzyme-A and luminescence was measured. Immediately after, an equal amount of Renilla Buffer 1X supplemented with 10 μM Coelenterazine was added to the sample and luminescence was measured again. Non-transfected lysate was used to measure the background signal, that was subtracted before calculating the firefly/renilla and renilla/firefly ratios.

### Flow cytometry analysis

Transfected cells were reseeded into 96-well tissue culture treated plates (Corning) at 30,000 cells/well containing varying doses of copper sulfate. Then, plates were incubated for 5 days to allow Cas13 expression and target KD. Finally, cells were subjected to automated flow cytometry using a high throughput autosampler connected to an LSR II (BD) at the Harvard Immunology Flow Core. Data was gated and quantified in BD software or Flow Jo software.

### Imaging analysis

Transfected cells were reseeded onto 384 Cell Carrier plates (Perkin Elmer) and imaged using an IN Cell Analyzer 6000 Cell Imaging System (GE) using a 20X air-immersion objective in widefield mode.

### Cloning

### UAS-Cas13b-NES-3XHA and UAS-dCas13b-NES-3XHA

The PspCas13b-NES-3XHA and dPspCas13b-NES-3XHA sequences were amplified by PCR from pC0046-EF1a-PspCas13b-NES-HIV and pC0049-EF1a-dPSPCas13b-NES-HIV, H133A/H1058A was a gift from Feng Zhang (Addgene plasmid # 103865; http://n2t.net/addgene:103865; RRID:Addgene_103865 and Addgene plasmid # 103862; http://n2t.net/addgene:103862; RRID:Addgene_103862) using the following primers: F: 5’-CACACCAGAATTCGCCACCATGAACATCCCCG-3’ R: 5’-ACACCACTCGAGTTAGGCATAGTCGGGGACAT-3’ The sequence was subcloned into pUASattB (Kadener’s Lab) using EcoRI and XhoI digestion.

### UAS-μS-Cas13b-NES-3XHA and UAS-μM-Cas13b-NES3X-HA

The μORFs sequences were amplified from the Firefly Cassette using the pMARL, a kind donation from Dr. Marr lab at Brandeis University, Waltham-MA. Specifically, the μS includes the first 22 aa of the coding sequence, while μM the first 38 aa. The primers used were μORF_Fire_F: 5’-gatctgcggccgcggcGCCACCATGGAAGACGCCAAAAAC-3’ μS_Fire_R: 5’-GAGCGGGGATGTTCATGTTATTAAGCGGTTCCATCCTCTAGAGG-3’ μM_Fire_R:GAGCGGGGATGTTCATGTTATTATCCAGGAACCAGGGCGTATC-3’ The PspCas13b-NES-3XHA sequence was amplified from the pC0046-EF1a-PspCas13b-NES-HIV with the following primers: R: 5’-CTAGAGGTACCCTCGATTAGGCATAGTCGGG-3’ μS_F: 5’-CCTCTAGAGGATGGAACCGCTTAATAACATGAACATCCCCGCTC-3’ μM_F: 5’-GATACGCCCTGGTTCCTGGATAATAACATGAACATCCCCGCTC-3’ The μORF and Cas13 fragments were assembled into pUASattB digested with XhoI by Gibson Assembly.

### UAS-NLS-Cas13b-NLS-3XHA

pC0046-EF1a-PspCas13b-NES-HIV was digested with NheI and EcoRI-HF, the removed sequence was replaced with a Gblock carrying part of Cas13b sequence linked to the 3XHA tag without the NES. The linker was changed from Gly-Ser-Ser (GGT-AGT-TCC) to Gly-Thre-Ser (GGT-ACC-TCC) to introduce a new KpnI site. To generate the UAS-Cas13b-3XHA, the Cas13b-3XHA sequence was subcloned into the pUASattB digested with XbaI and EcoRI using the following primers: F: 5’-CACACCAGAATTCGCCACCATGAACATCCCCG-3’ R: 5’-CACCATCTAGATTAGGCATAGTCGGGGACATCAT To generate *UAS-Cas13b-NLS-3XHA, UAS-Cas13b-3XHA* was digested with KpnI and ligated to the following pre-annealed oligos: Top_NLS_Ct: 5’-CTCCCCTAAGAAAAAGAGGAAGGTGGGTAC-3’ Bottom_NLS_Ct:5’-CCACCTTCCTCTTTTTCTTAGGGGAGGTAC-3’ Finally, to generate the *UAS-NLS-Cas13b-NLS-HA* plasmid, *UAS-Cas13b-NLS-HA* was amplified with the following primers: F:5’CACCAAATTCGCCACCATGGGTACCTCCCCTAAGAAAAAGAGGAAGAACATCCCCGCTCTGGTG-3’ R:5’-CACCATCTAGATTAGGCATAGTCGGGGACATCAT And inserted into the pUAS-attB digested with EcoRI and XbaI.

### UAS-Cas13d-NES-3XHA

Cas13d was amplified from pXR001: EF1a-CasRx-2A-EGFP (Addgene # #109049) using the primers 5’ caagaagagaactctgaatagggaattgggaattcgccaccATGatcgaaaaaaaaaagtccttcgccaagggc-3’ and 5’-ggaattgccggacaccttctttttctcctt-3.

NES-3XHA was amplified from UAS-Cas13d-NES-3XHAusing 5’ aaggagaaaaagaaggtgtccggcaattccggcagcctgcagctgcccccgctggagcgc-3’ and 5’-tctctgtaggtagtttgtccaattatgtca-3’, and then the two fragments were inserted between the EcoRI and XhoI sites of pUASattB-PspCas13b using Gibson Assemly.

### UAS-NLS-CasRx-NLS-3XHA

NLS-Cas13d-NLS was amplified from pXR001: EF1a-CasRx-2A-EGFP (Addgene # #109049) using primers: 5’caagaagagaactctgaatagggaattgggaattcgccaccatgagccccaagaagaagagaaaggtggaggcc agc-3’ and 5’-tcgtaggggtaggcgtaatctggcacgtcgtacgggtaagcggccgccaccttcctctttttcttagg -3’ The3XHA was amplified using 5’-cctaagaaaaagaggaaggtggcggccgcttacccatacgatgttccagattacgct-3’ and 5’-tctctgtaggtagtttgtccaattatgtca-3’, and then the two fragments were inserted between the EcoRI and XhoI sites of pUASattB-PspCas13b using Gibson Assemly.

#### crRNA plasmids

To create the pU6.3crRNA-Cas13b plasmid, we digested pCFD3.1-w-dU6:3gRNA, a gift from Simon Bullock (Addgene plasmid # 123366; http://n2t.net/addgene:123366; RRID:Addgene_123366), with BbsI and XbaI and ligated it to a Gblock carrying the crRNA scaffold for Cas13b. To introduce the protospacer, we pre annealed oligos and inserted into the pU6.3crRNA-Cas13b plasmid by Golden Gate with BbsI. The oligos used for the cloning are the following:

renilla_top_U63: gtcgGATCAAGTAACCTATAAGAACCATTACCAG

renilla_bot_U63: caacCTGGTAATGGTTCTTATAGGTTACTTGATC

firefly_top_U6.3 gtcgGACACATAATTCGCCTCTCTGATTAACGCC

firefly_bot_U6.3:caacGGCGTTAATCAGAGAGGCGAATTATGTGTC

cherry_top_U63: gctgGGGAGGTGATGTCCAACTTGATGTTGACG

cherry_bot_U63: caacCGTCAACATCAAGTTGGACATCACCTCCC

To create the crRNA vectors that allowed co-expression of GFP or EBFP2, U6:3 promoter, guide acceptor region, and tracrRNA was amplified from pCFD3-dU6:3gRNA (Addgene # 49410) and inserted into pLib6.4 (Addgene # #133783) between BstBI and KpnI to generate pLib6.6. EBFP2 was amplified from pEBFP2-Nuc (Addgene # 14893) and inserted into pLib vectors between PmeI and NheI to generate pLib6.4B or pLib6.6B. To introduce the RfxCas13d direct repeat (which is 5’ to the protospacer), pLib6.6 or pLib6.6B were digested with BbsI and annealed oligos were introduced which recreated the paired BbsI sites following the direct repeat creating pLib11.6 and pLib11.6B, allowing direct cloning of crRNAs with the 5’ addition of 5’-AAAC-3’ to the sense primer and the 5’ addition of 5’-AAAA-3’ to the antisense primer. To introduce the PspCas13b direct repeat (which is 3’ to the protospacer), pLib6.6 or pLib6.6B were digested with BbsI and annealed oligos were introduced which recreated the paired BbsI sites creating pLib11.7 and pLib11.7B, allowing direct cloning of crRNAs with the 5’ addition of 5’-GTCG-3’ to the sense primer and the 5’ addition of 5’-CAAC-3’ to the antisense primer. In all experiments reported here, pLib11.6B and pLib11.7 were used for cloning crRNAs. The sequences were as follows:

mCherry_inv: GGAGGATAACATGGCCATCATCAAGGAGT

mCherry_1: ACTCCTTGATGATGGCCATGTTATCCTCC

mCherry_2: GGGAGGTGATGTCCAACTTGATGTTGACG

mCherry_3: AGTCCTCGTTGTGGGAGGTGATGTCCAAC

mCherry_4: CTTCAGCTTCAGCCTCTGCTTGATCTCG

Rho1_7: GATCTTTGCTGAAGACAATCAGAAGGCAA

Rho1_8: ACGTCAGTGTCGGGATAGCTCAGCGGTCG

Rho1_9: CCTACCAAAATGATTGGAACATTTGGAC

Rho1_10: GACCCTCCTGCGGCTTCACCGGCTCCTGCT

### MT-NLS-Cas13d-NLS-HA-T2A-GFP

RfA Gateway® cassette (Thermo) was inserted into pMK33 (PMID: 1913820) between XhoI and SpeI sites to generate pMK33-GW. NLS-Cas13d-NLS-HA-T2A-GFP was amplified from pXR001: EF1a-CasRx-2A-EGFP (Addgene #109049) and recombined into pCR8-TOPO and then further recombined into pMK33-GW to generate pMK33/NLS-Cas13d-NLS-HA-T2A-GFP. pMK33/NLS-Cas13d-NLS-HA-T2A-GFP was transfected into S2R+-NPT005 cells (DGRC # 229) using Effectene (Qiagen) according to manufacturer’s protocol and selected in 200 ng/mL Hygromycin B (Calbiochem) for one month. Then, surviving cells were further subcloned by transferring individual GFP-negative cells to 96-well plates by Fluorescence-activated cell sorting using a FACSAria (BD). 20 clones were expanded and visually reassessed for GFP-inducibility in the presence of 100 μM CμSO4. This identified one clone (F11) with strictly inducible GFP expression, which also strictly induced expression of NLS-Cas13d-NLS-HA, verified by HA Western blot (data not shown), and was used for all subsequent experiments. Clone D7, with constitutive activity, was used in Supplementary Figure 3.

## References

1. Brand, A. H. & Dormand, E. The GAL4 system as a tool for unravelling the mysteries of the. 572–578 (1995).

2. Perkins, L. A. et al. The Transgenic RNAi Project at Harvard Medical School?: Resources and Validation. Genetics 201, 843–852 (2015).

3. Housden, B. E. & Perrimon, N. Comparing CRISPR and RNAi-based screening technologies. Nat. Biotechnol. 34, 621–623 (2016).

4. Jr, W. G. K. Use and Abuse of RNAi to Study. Science (80-.). 337, 421–423 (2012).

5. Makarova, K. S., Grishin, N. V, Shabalina, S. A., Wolf, Y. I. & Koonin, E. V. A putative RNA-interference-based immune system in prokaryotes?: computational analysis of the predicted enzymatic machinery, functional analogies with eukaryotic RNAi, and hypothetical mechanisms of action. Biol. Direct 26, 1–26 (2006).

6. Ophir Shalem, Neville E. Sanjana, Ella Hartenian, Xi Shi, David A. Scott, Tarjei S. Mikkelsen, Dirk Heckl, Benjamin L. Ebert, David E. Root, John G. Doench, F. Z. Genome-Scale CRISPR-Cas9 Knockout Screening in Human Cells. Science (80-.). 343, 84–88 (2014).

7. Yang, L. et al. RNA-Guided Human Genome Engineering via Cas9. Science (80-.). 823–827 (2013).

8. Le Cong, F. Ann Ran, David Cox, Shuailiang Lin, Robert Barretto, Naomi Habib, Patrick D. Hsu, Xuebing Wu, 7 Wenyan Jiang, Luciano A. Marraffini, F. Z. Multiplex Genome Engineering Using CRISPR/Cas Systems. Science (80-.). 339, 819–824 (2013).

9. Jia, Y. et al./ Next-generation CRISPR/Cas9 transcriptional activation in Drosophila using flySAM. PNAS 115, 4719–4724 (2018).

10. Qi, L. S. et al. Repurposing CRISPR as an RNA-guided platform for sequence-specific control of gene expression. Cell 152, 1173–1183 (2013).

11. Gilbert, L. A. et al. CRISPR-mediated modular RNA-guided regulation of transcription in eukaryotes. Cell 154, 442–451 (2013).

12. Gootenberg, J. S. et al. Nucleic acid detection with CRISPR-Cas13a/C2c2. Science (80-.). 442, 438–442 (2017).

13. Abudayyeh, O. O. et al. RNA targeting with CRISPR–Cas13. Nature 550, 280–284 (2017).

14. Cox, D. B. T., Gootenberg, J. S. & Abudayyeh, O. O. RNA editing with CRISPR-Cas13. Science (80-.). 1027, 1019–1027 (2017).

15. Winston X. Yan, Shaorong Chong, Huaibin Zhang, Kira S. Makarova, Eugene V. Koonin, David R. Cheng, D. A. S. Cas13d Is a Compact RNA-Targeting Type VI CRISPR Effector Positively Modulated by a WYL-Domain-Article Cas13d Is a Compact RNA-Targeting Type VI CRISPR Effector Positively Modulated by a WYL-Domain-Containing Accessory Protein. Mol. Cell 70, 327–339 (2018).

16. Connell, M. R. O. Molecular Mechanisms of RNA Targeting by Type VI CRISPR – Cas Systems. J. Mol. Biol. 431, 66–87 (2019).

17. East-seletsky, A. et al. Two distinct RNase activities of CRISPR-C2c2 enable guide-RNA processing and RNA detection. Nature 538, 270–273 (2016).

18. Bandaru, S. et al. Structure - based design of gRNA for Cas13. Sci. Rep. 10, 1–12 (2020).

19. Silvana Konermann, Peter Lotfy, Nicholas J. Brideau, Jennifer Oki, Maxim N. Shokhirev, P. D. H. Transcriptome Engineering with RNA-Targeting Type VI-D CRISPR Effectors Graphical. Cell 173, 665-668.e14 (2018).

20. Gootenberg, J. S. et al. Multiplexed and portable nucleic acid detection platform with Cas13, Cas12a, and Csm6. Science (80-.). 444, 439–444 (2018).

21. Kushawah, G. et al. Technology CRISPR-Cas13d Induces Efficient mRNA Knockdown in Animal Embryos ll Technology CRISPR-Cas13d Induces Efficient mRNA Knockdown in Animal Embryos. Dev. Cell 54, 805–817 (2020).

22. Buchman, A. B. et al. Programmable RNA Targeting Using CasRx in Flies. Cris. J. 3, 164–176 (2020).

23. Neumüller, R. A. et al. Stringent Analysis of Gene Function and Protein-Protein Interactions Using Fluorescently Tagged Genes. Genetics 190, 931–940 (2012).

24. Port, F. et al. A large-scale resource for tissue-specific CRISPR mutagenesis in Drosophila. Elife 9, 1–20 (2020).

25. Prokopenko, S. N. et al. A putative exchange factor for Rho1 GTPase is required for initiation of cytokinesis in Drosophila. Genesis 13, 2301–2314 (1999).

26. Magie, C. R., Pinto-santini, D. & Parkhurst, S. M. Rho1 interacts with p120ctn and α-catenin, and regulates cadherin-based adherens adherens junction components in Drosophila. Develop 3782, 3771–3782 (2002).

27. Rogers, S. L. & Rogers, G. C. Culture of Drosophila S2 cells and their use for RNAi-mediated loss-of-function studies and immunofluorescence microscopy. Nat. Protoc. 3, 606–611 (2008).

28. Wang, H. et al. CRISPR-mediated live imaging of genome editing and transcription. Science (80-.). 365, 1301–1305 (2019).

29. Wilson, C., Chen, P. J., Miao, Z. & Liu, D. R. Programmable m6A modification of cellular RNAs with a Cas13-directed methyltransferase. Nat. Biotechnol. June, (2020).

30. Li, J. et al. Targeted mRNA demethylation using an engineered dCas13b-ALKBH5 fusion protein. Nucleic Acids Res. 48, 5684–5694 (2020).

31. Menghan Du, Nathaniel Jillette, Jacqueline Jufen Zhu, Sheng Li, A.W.C. CRISPR artificial splicing factors. Nat. Commun. 11, 1–11 (2020).

32. Housden, B. E. et al. Improved detection of synthetic lethal interactions in Drosophila cells using variable dose analysis (VDA). PNAS November, 10755–10762 (2017).

33. Viswanatha, R., Li, Z., Hu, Y. & Perrimon, N. Pooled genome-wide CRISPR screening for basal and context-specific fitness gene essentiality in Drosophila cells. Elife 7, (2018).

