## Supplementary Figures for "CRISPR-Cas13 mediated Knock Down in *Drosophila* cultured cells"

### Supplementary Figure 1

## A

Gate all cells on EBFP intensity (P6)

↓

Measure EBFP intensity in P6 (report in Figure 1G)

↓

Gate P6 cells on mCherry intensity (P7) (report P7 as a fraction of P6 in Figure 1F)

## B

MT-NLS-Cas13d-NLS-HA-T2A-GFP
+ 10 µM copper sulfate
+ crRNA [mCherry_inv]::EBFP

MT-NLS-Cas13d-NLS-HA-T2A-GFP
+ 10 µM copper sulfate
+ crRNA [mCherry_3]::EBFP

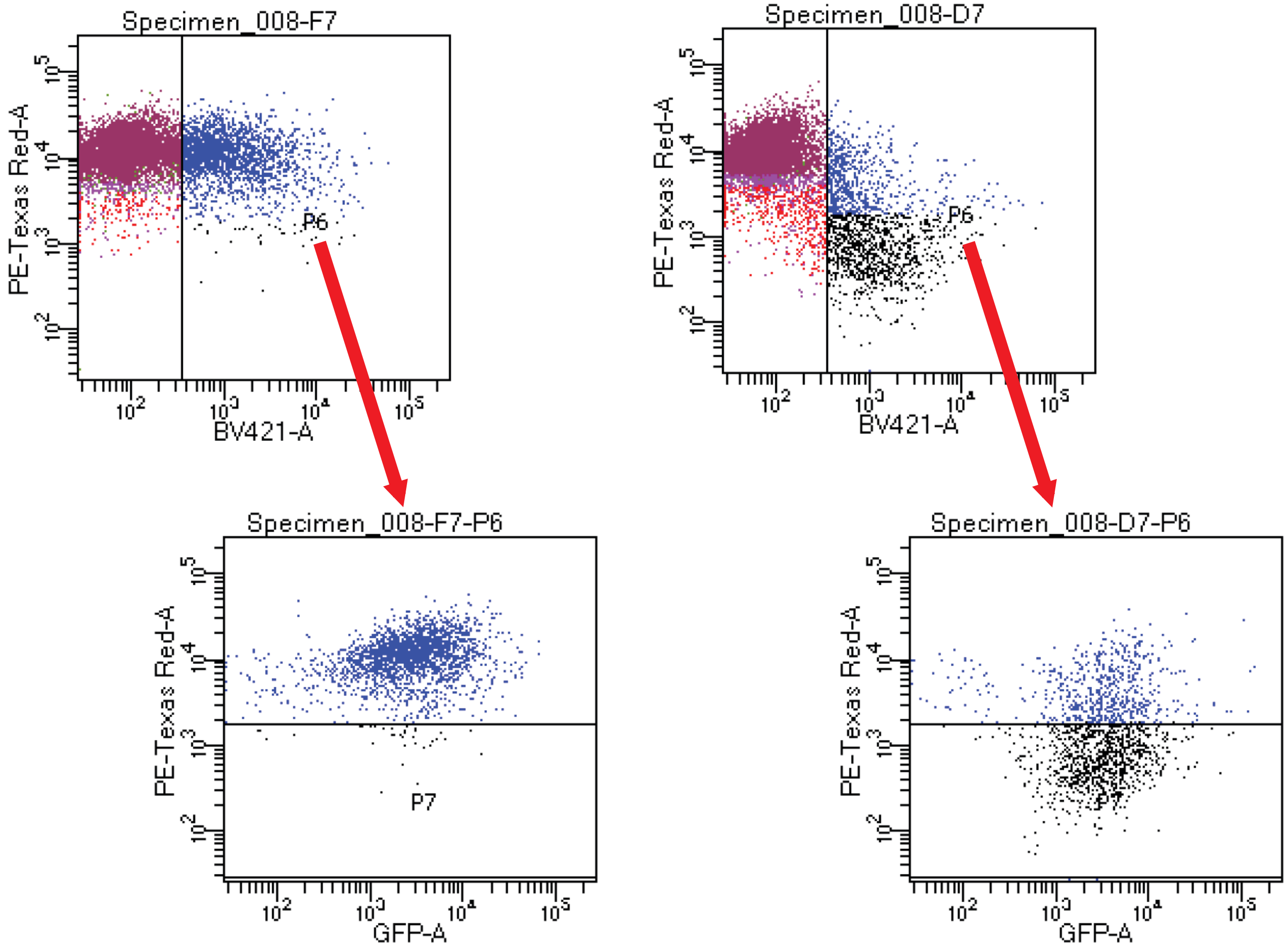

### Supplementary Figure 2

A

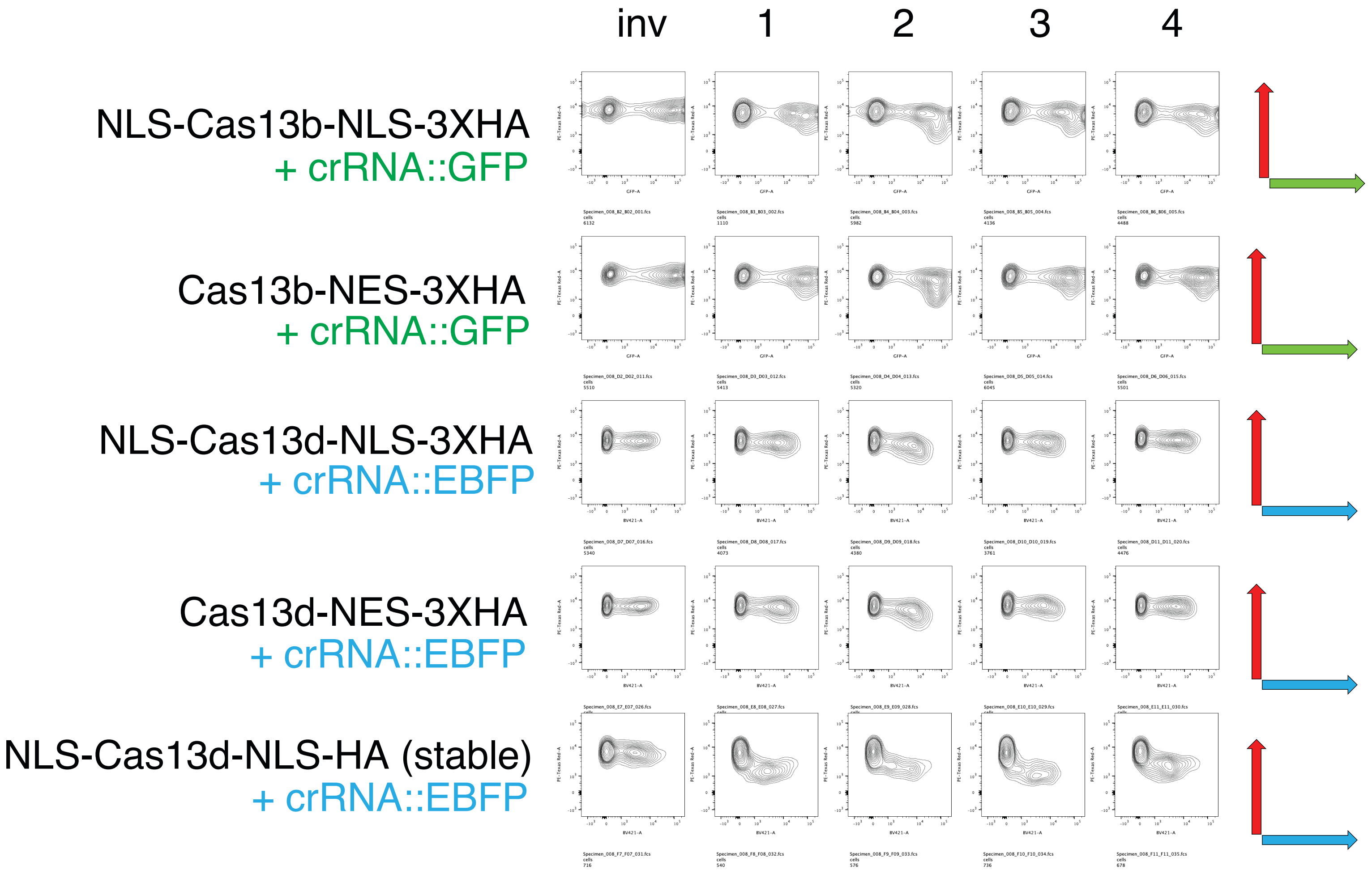


B

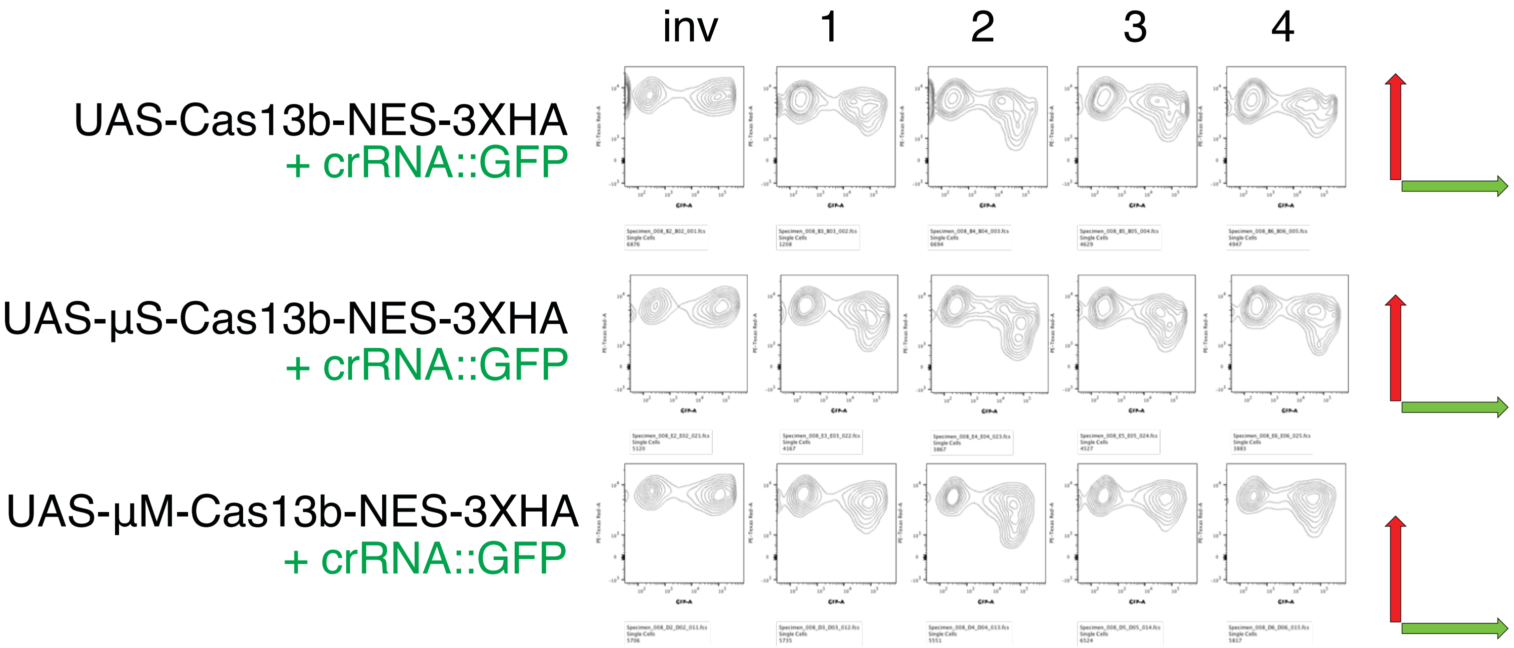

### Supplementary Figure 3

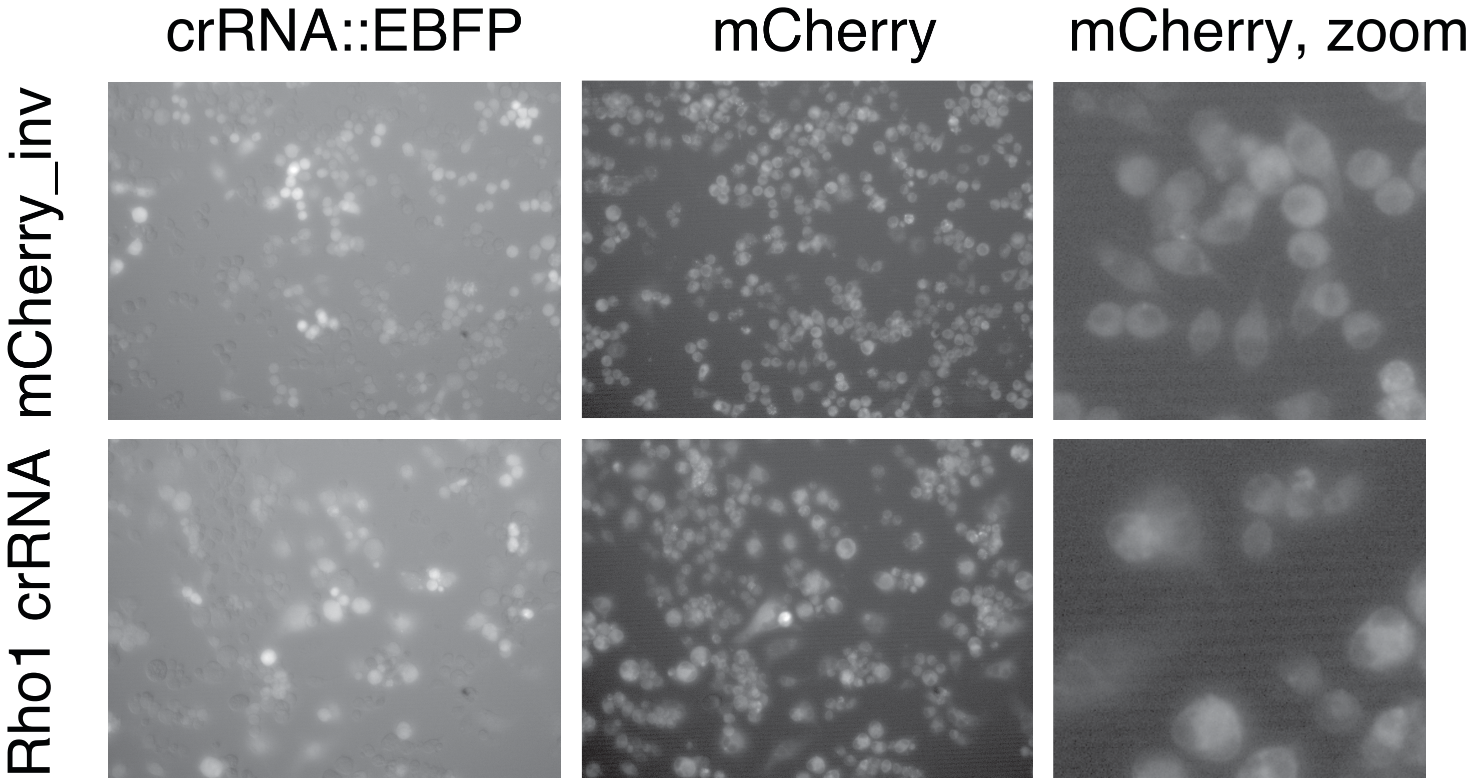
